# Chromatin structure and 3D architecture define differential functions of *PU.1* cis regulatory elements in human blood cell lineages

**DOI:** 10.1101/2024.01.01.573782

**Authors:** Kevin Qiu, Duc Vu, Leran Wang, Anna Bookstaver, Thang N. Dinh, Adam N. Goldfarb, Daniel G. Tenen, Bon Q. Trinh

**Author notes:** Corresponding Author and Lead Contact: Bon Q. Trinh.

## Abstract

The precise spatio-temporal expression of the hematopoietic ETS transcription factor *PU.1* that determines the hematopoietic cell fates is tightly regulated at the chromatin level. However, it remains elusive as to how chromatin signatures are linked to this dynamic expression pattern *of PU.1* across blood cell lineages. Here we performed an unbiased and in-depth analysis of the relationship between human *PU.1* expression, the presence of trans-acting factors, and 3D architecture at various cis-regulatory elements (CRE) proximal to the *PU.1* locus. We identified multiple novel CREs at the upstream region of the gene following an integrative inspection for conserved DNA elements at the chromatin-accessible regions in primary human blood lineages. We showed that a subset of CREs localize within a 10 kb-wide cluster that exhibits that exhibit molecular features of a myeloid-specific super-enhancer involved in mediating *PU.1* autoregulation, including open chromatin, unmethylated DNA, histone enhancer marks, transcription of enhancer RNAs, and occupancy of the PU.1 protein itself. Importantly, we revealed the presence of common 35-kb-wide CTCF-bound insulated neighborhood that contains the CRE cluster, forming the chromatin territory for lineage-specific and CRE-mediated chromatin interactions. These include functional CRE-promoter interactions in myeloid and B cells but not in erythroid and T cells. Our findings also provide mechanistic insights into the interplay between dynamic chromatin structure and 3D architecture in defining certain CREs as enhancers or silencers in chromatin regulation of *PU.1* expression. The study lays the groundwork for further examination of *PU.1* CREs as well as epigenetic regulation in malignant hematopoiesis.

## INTRODUCTION

Changes to the accessibility, methylation, histone post-translational modifications, and 3D architectures of chromatin structures in eukaryotes are known to determine the expression patterns of lineage-defining genes (Gibney and Nolan 2010). In blood, the precise expression of the ETS-family transcription factor PU.1 (also known as SPI1) determine the differentiation journeys undertaken by various hematopoietic cells in the bone marrow. *PU.1* expression is both highest in and absolutely necessary for the development of myelomonocytic cells (Klemsz, McKercher et al. 1990, Scott, Simon et al. 1994, McKercher, Torbett et al. 1996, DeKoter and Singh 2000, Iwasaki, Somoza et al. 2005, Beers, Henkel et al. 2006, Rosenbauer, Owens et al. 2006). Lower expression is needed for B cells (Polli, Dakic et al. 2005), and residual *PU.1* expression is required for the proliferation of erythroid progenitors (Choe, Ujhelly et al. 2010, Pop, Shearstone et al. 2010, Wontakal, Guo et al. 2011). In the T cell lineage, *PU.1* expression is reduced in a stepwise manner beginning in progenitors and is fully extinguished in mature cells (Rosenbauer, Owens et al. 2006). Deviations from this natural expression pattern are associated with multiple blood diseases, including malignancy (Chen, Zhang et al. 1995, Rosenbauer, Owens et al. 2006). Although some studies have revealed the association of histone marks with *PU.1* expression in myeloid cells (Hoogenkamp, Krysinska et al. 2007, Trinh, Ummarino et al. 2021), it is not fully understood as to what and how chromatin signatures are linked to the dynamic expression pattern *of PU.1* among blood cell lineages.

Studies over the past two decades have demonstrated the role of cis-regulatory elements (CRE) in regulating *PU.1* expression. The CREs act through long distance and within the chromatin context. The highly conserved and well-studied upstream regulatory element (URE) (human -17.2 kb, murine -14 kb) induces *PU.1* expression in myeloid, B, and erythroid progenitors (Li, Okuno et al. 2001, Rosenbauer, Wagner et al. 2004, Okuno, Huang et al. 2005, Rosenbauer, Owens et al. 2006, Bonadies, Neururer et al. 2010, Will, Vogler et al. 2015, Willcockson, Taylor et al. 2019). In contrast, it represses *PU.1* expression in progenitor T cells (Rosenbauer, Owens et al. 2006). In addition to the URE, several less-known regulatory elements have been described in the murine genome. For instance, the ZL12 element (stands for Leddin & Zarnegar -murine 12 kb) was reported to synergize with the URE to mediate high murine *Pu.1* expression exclusively in mature myeloid cells (Zarnegar, Chen et al. 2010, Leddin, Perrod et al. 2011). Another CRE called CE5 induces transient *Pu.1* promoter activity in murine macrophages (Zarnegar, Chen et al. 2010). On the other hand, the CE4A (murine -9 kb) inhibits *Pu.1* promoter activity in the context of chromatin in a murine T cell line (Zarnegar, Chen et al. 2010). The identification of functional PU.1 CREs in mouse models warrants a comprehensive survey of CREs in human hematopoietic cells in order to better understand the regulation of physiologic expression patterns of human *PU.1*.

There is increasing evidence indicating that *PU.1* CREs can function as enhancers, silencers, or both, depending on the blood cell lineage and the nature of the element itself. It is well documented that the same CRE can function as an enhancer or silencer depending on cellular conditions and types as well as tissue contexts (Simpson, Schell et al. 1986, Jiang, Cai et al. 1993, Bessis, Champtiaux et al. 1997, Kallunki, Edelman et al. 1998, Kehayova, Monahan et al. 2011, Gisselbrecht, Palagi et al. 2020). Common features between enhancers and silencers have been implicated as having similar physical contacts with target promoters and localizing in chromatin accessible regions (Huang, Petrykowska et al. 2019). Moreover, post-translational histone modifications have been increasingly utilized to infer the epigenetic characteristics of CREs across cellular contexts. For instance, active enhancers are generally marked with Histone 3 lysine 27 acetylation (H3K27Ac) (Creyghton, Cheng et al. 2010, Pekowska, Benoukraf et al. 2011). Histone 3 lysine 4 mono-methylation (H3K4me1) and H3 lysine 4 di-methylation (H3K4me2) are considered as indicators for the presence of enhancers and gene transcription (Creyghton, Cheng et al. 2010, Pekowska, Benoukraf et al. 2011). On the other hand, implicated silencers can be identified by the presence of the histone 3 lysine 27 tri-methylation (H3K27Me3) repressive mark (Young, Willson et al. 2011, Huang, Petrykowska et al. 2019) and the H3K9Me3 heterochromatin mark (Nicetto, Donahue et al. 2019). One example of a bifunctional *PU.1* CRE is the URE, which was characterized as an enhancer in myeloid cells but a silencer in T cells (Rosenbauer, Wagner et al. 2004, Rosenbauer, Owens et al. 2006, Will, Vogler et al. 2015, Willcockson, Taylor et al. 2019). Indeed, the function of the URE as an enhancer in myeloid cells has been well documented in both human and mouse models through the enrichment of H3K27Ac, H3K4Me1, and H3K4Me2 chromatin marks, and promoter contact (Li, Okuno et al. 2001, Rosenbauer, Wagner et al. 2004, Okuno, Huang et al. 2005, Rosenbauer, Owens et al. 2006, Bonadies, Neururer et al. 2010, Will, Vogler et al. 2015, Trinh, Ummarino et al. 2021), but the molecular signatures for URE and the identification of other potential *PU.1* silencers have not been investigated. Overall, a comprehensive investigation of chromatin signatures at the *PU.1* locus is needed.

In this study, we performed a comprehensive investigation of *PU.1* CREs and their chromatin signatures throughout the human *PU.1* locus in human primary blood cells. Beginning with an integrative survey for conserved DNA elements at the chromatin accessible regions, we identified an 8-kb CRE cluster containing the URE and several unknown CREs. The cluster exhibits myeloid-specific enhancer features, including chromatin opening, DNA unmethylation, H3K27Ac enrichment, the presence of enhancer RNAs, and the occupancy of *PU.1,* which mediates autoregulation. Although the presence of *PU.1* silencers in T cells has been documented (Rosenbauer, Wagner et al. 2004, Rosenbauer, Owens et al. 2006, Zarnegar, Chen et al. 2010, Will, Vogler et al. 2015, Willcockson, Taylor et al. 2019), we noted that the repressive mark H3K27Me3 and the heterochromatin mark H3K9Me3 were not specifically demarcated for *PU.1* silencers. Instead, we revealed the presence of unique and dynamic 3D chromatin architectures at the *PU.1* locus. Our findings provide mechanistic insights into the role of epigenetic and chromatin regulation of *PU.1* and lay the groundwork for further examination of individual CRE constituent in normal and malignant hematopoiesis.

## METHODS

### ATAC-seq data analyses

Raw ATAC-Seq fastq files were downloaded from GSE74912 using SRA-Toolkit. Read quality was assessed by FastQC (Andrews 2010). Fastq files were adapter and quality trimmed using Trim-Galore using default settings and aligned to hg38 using bowtie2 (Langmead and Salzberg 2012). The aligned reads were then filtered for unmapped and mitochondrial reads as well as PCR duplicates using Samtools (Danecek, Bonfield et al. 2021) and Picard Tools (https://broadinstitute.github.io/picard/). Filtered reads on the positive and negative strands were shifted +4bp and -5bp respectively to adjust for Tn5 transposase binding and reads aligning to blacklisted regions were removed using BedTools (Quinlan and Hall 2010). Peaks were then called using MACS2 using parameters “-f BED -g hs -q 0.01”. Counts for ATAC-Seq consensus peaks as well as regions of interest were generated using DiffBind, and scaling factors for each sample were calculated using EdgeR’s TMM normalization function (Robinson, McCarthy et al. 2010, Stark and Brown 2023). The reciprocal size factors were then used to generate normalized BigWig files using DeepTool’s bamCoverage function.

### ChIP-seq data analyses

Transcription factor and histone ChIP-Seq datasets from ENCODE and GEO were downloaded and adapter-trimmed as previously described in the ATAC-Seq methods and aligned to hg38 using the ENCODE ChIP-Seq Uniform Processing Pipeline v1.4 (Hitz, Lee et al. 2023), with peaks being called by MACS2 (Zhang, Liu et al. 2008). Filtered bam and peak files were then used to generate normalized BigWig files, binding plots, and counts using DiffBind.

### PRO/GRO-seq data analyses

PRO/GRO-seq data and GRO-seq fastqs were downloaded from GEO and adapter sequences were identified using FastQC. Fastqs were aligned to hg38 using the Proseq2.0 pipeline using the adaptor with the “-4DREG” and “—ADAPT/--ADAPT1/--ADAPT2” parameters. Plus and minus stranded BigWigs were run through the DREG peak calling pipeline. Filtered bams and peak files were then used to generate binding plots and counts.

### WGBS data analyses

Whole-genome bisulfite sequencing data .pat and .beta files were downloaded GSE186458. Methylated probes for regions of interest were binned and normalized to proportion of probes methylated using wgbs_tools (Loyfer, Magenheim et al. 2023) and plotted in a heatmap using ComplexHeatmap R package.

### Hi-C data analyses

Processed MNase Hi-C dataset files (.bam, .hic, and .pairs aligned to hg38) were downloaded from ENCODE. GEO MNase datasets for primary erythroid cells were downloaded from GSE214808 (Li, Zhao et al. 2023) aligned to hg38 and processed via the ENCODE HI-C Uniform Processing Pipeline v1.15.1. Chromatin loops were called using using Juicer’s HiCCUPS at a 1kb resolution, and HiC matrices were plotted using Juicebox (Durand, Shamim et al. 2016).

### Histone mark browser track visualization

BigWig averages were calculated using Wiggletools, and binding plots of regions of interest were created using DeepTool’s computeMatrix and plotProfile functions. To visualize the histone marks via tracks, the online program UCSC genome browser was used to upload the bigwig files on. Vertical viewing range varies with overall peak intensity.

### RNA sequencing data analysis (RNA-seq)

Cell line bulk RNA-Seq datasets were downloaded, and adapter trimmed as previously described. Filtered fastqs were then aligned to hg38 using STAR 2.7.9a (Dobin, Davis et al. 2013), and counts files were generated using htseq-count 0.13.5. Counts were normalized using DESeq2 (Love, Huber et al. 2014).

### Single-cell RNA-seq (scRNA-seq) data analyses

Single Cell Multiome ATAC and Gene Expression datasets processed by Cell Ranger ARC 2.0.0 for mononuclear cells isolated from peripheral blood of a healthy donor were obtained from the 10x Genomics public datasets repository and analyzed using Signac and Seurat (McGinnis, Murrow et al. 2019, Stuart, Srivastava et al. 2021). Predicted doublets were identified and removed using doubletFinder, and ells containing less than 1000 RNA/ATAC counts, >2 nucleosome signal score, <TSS enrichment score, <0.25 fraction reads in peak, and greater than 25 percent mitochondrial RNA were filtered using Seurat’s subset function. Cells were annotated based on a previously annotated dataset (GSE181061) using Seurat’s FindTransferAnchors function. Cells identified as myeloid (monocytic), B, and T cells (CD4/CD8/Tregs), and were plotted using a multimodal UMAP generated using FindMultiModalNeighbors.

## RESULTS

### Identification of putative cis-regulatory elements throughout the *PU.1* locus

We first assessed the relationship between sequence conservation across species and chromatin accessibility across the *PU.1* gene locus. By interrogating bulk ATAC-seq data from four primary human blood cell lineages (Corces, Buenrostro et al. 2016) and the Cons 100 Verts track with multiple alignments of 100 vertebrate species and measurements of evolutionary conservation (http://genome.ucsc.edu), we identified 14 DNA regions that exhibit sequence conservation and chromatin opening in at least one in four major human definitive blood cell lineages (Figures 1A-1B). Among these are the URE, which contains two conserved regions and has been extensively studied in human and murine models (Li, Okuno et al. 2001, Okuno, Huang et al. 2005, Bonadies, Neururer et al. 2010), the core *PU.1* promoter (PrPr), and the and the antisense promoter (AsPr) for a *PU.1* antisense lncRNA (Ebralidze, Guibal et al. 2008, van der Kouwe, Heller et al. 2021). We also identified human homologs two CREs previously reported in murine models, the ZL12 (Zarnegar, Chen et al. 2010, Leddin, Perrod et al. 2011) and the CE5 (Zarnegar, Chen et al. 2010), which we will refer to as hZL12 and hCE5. Moreover, we identified eight putative CREs that we called TL1-8 (Figure 1A, lower panel). Among the four major human blood cell lines, chromatin accessibility is generally superior in myeloid cells across nearly all the previously defined and novel CREs (TL5, hZL12, TL7, hCE5, TL8), although TL1 is less lineage specific (Figure 1C). The presence of ten uncharacterized DNA elements in human exhibiting sequence conservation and chromatin accessibility is suggestive of their potential involvement in lineage-specific *PU.1* expression.

**Figure 1.**
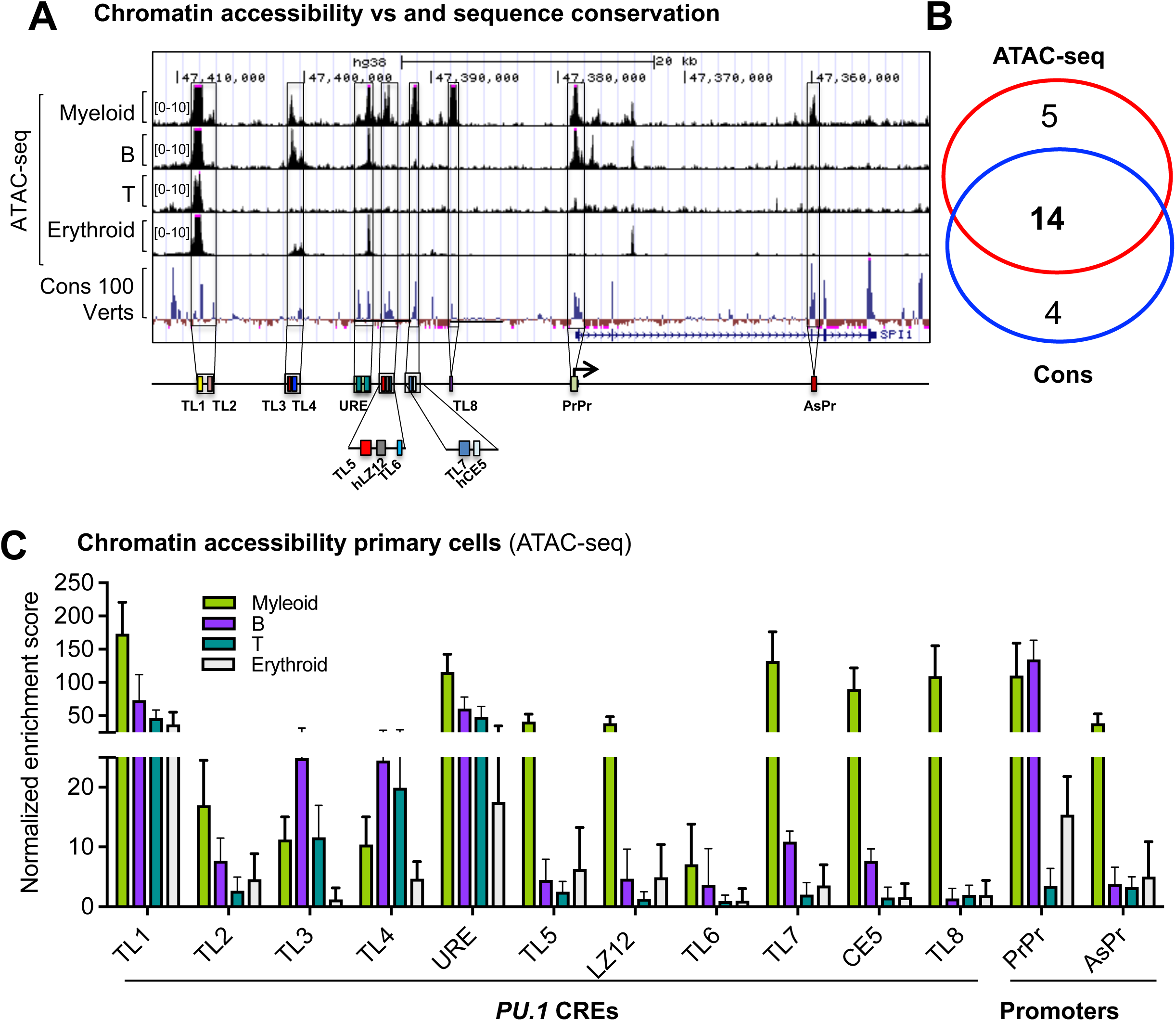
Conserved DNA elements with chromatin opening at the *PU.1* locus. A) Genomic track view of the genomic region covering the *PU.1* locus and its upstream region. Shown are ATAC-seq tracks of four human blood cell lineages (data was extracted from a published dataset (GEO: GSE74912) (upper panels), 100 vertebrates Basewise Conservation by PhyloP (middle panel), and diagram of DNA elements. B) Venn diagram showing DNA elements exhibiting sequence conservation and chromatin opening. C) Bar graph of ATAC-seq enrichment scores at the DNA elements from four human blood cell lineages.

### Chromatin accessibility at certain CREs is correlated with the unique expression pattern of *PU.1* in blood cell lineages

We next investigated the relationship between chromatin accessibility at the CREs and the expression pattern of *PU.1* in blood cell lineages. Using cellular indexing of transcriptomes and epitopes by sequencing (CITE-seq), which integrates cellular protein and transcriptome measurements at single-cell resolution (Stoeckius, Hafemeister et al. 2017), we verified the *PU.1* expression pattern in blood cell lineages. *PU.1* expression was highest in primary myeloid cells, less in B cells, minimally in erythroid cells, and essentially silenced in mature T cells (Figures 2A-B). This stepwise reduction in *PU.1* expression represents an important functional readout as we interrogated the role of CREs and epigenetic features in the regulation of *PU.1* expression. We next performed a multiomic analysis of a human peripheral blood mononuclear cell (PBMC) dataset (10x Genomics), which contains chromatin accessibility data and gene expression data within the same cell. Of note, chromatin accessibility at several elements, including URE, TL5, hLZ12, TL7, hCE5, and TL8, was specific to myeloid cells (Figure 2C, top panel). TL1 was highly accessible in but not lineage-specific, whereas TL3 and TL4 were more accessible in B and T cells (Figure 2C). In line with modest *PU.1* expression in B cells, chromatin accessibility was less at the URE but comparable at the PrPr (Figure 2C, middle panel). As *PU.1* expression is typically muted in T cells, chromatin was found to be inaccessible at the PrPr and most of the CREs, except for weak accessibility at the URE (Figures 2C, bottom panel). Overall, the differential chromatin accessibility at the CREs correlated well with lineage-specific differences in *PU.1* expression, underscoring the importance of further investigating the contribution of epigenetic signatures.

**Figure 2.**
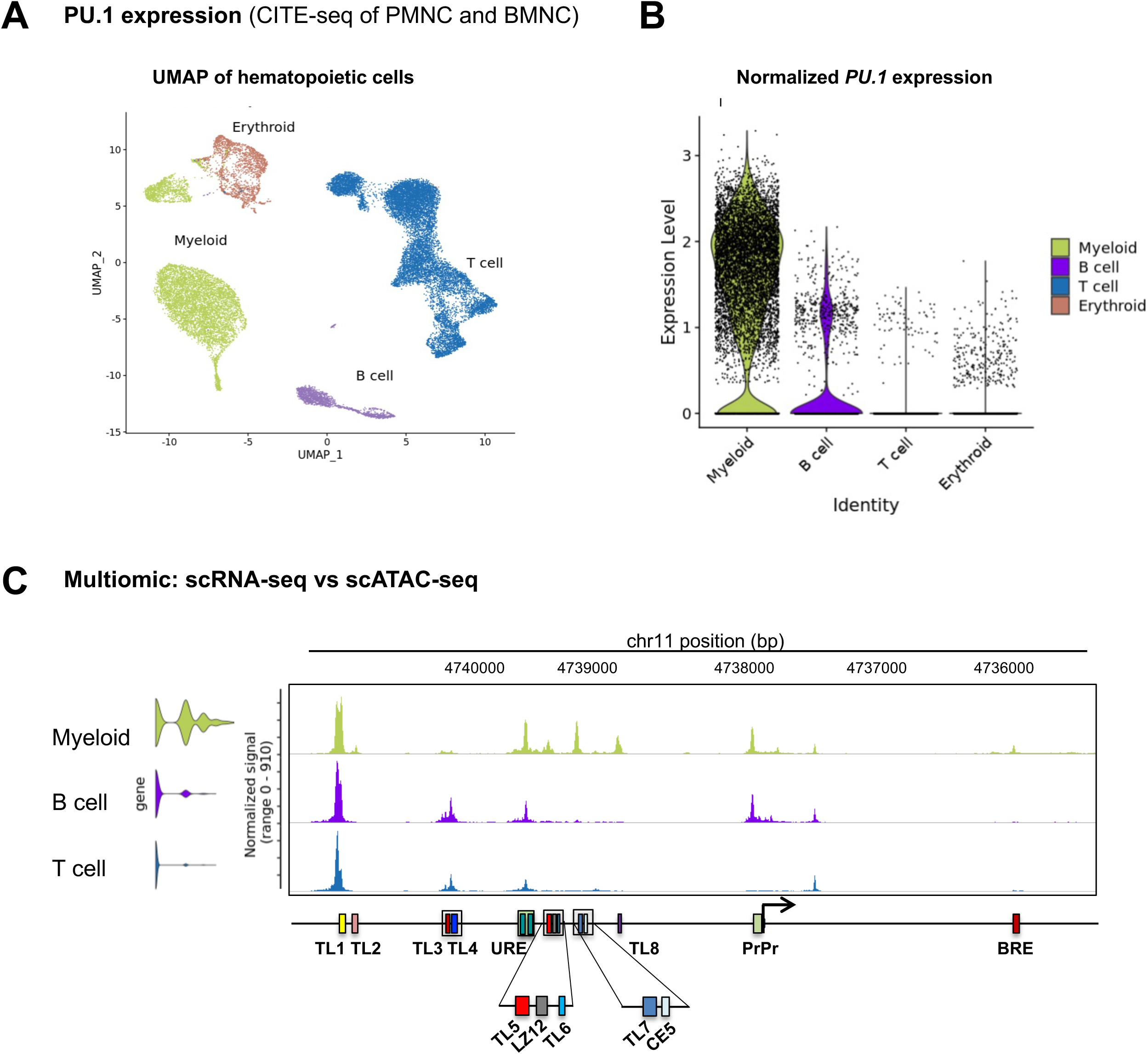
The association between *PU.1* expression and chromatin accessibility at certain regulatory elements. A-B) Expression analysis of *PU.1* in primary human blood cells at single-cell resolution. A) UMAP plot of myeloid, B, T, and erythroid cells. B) Transcript profiles of *PU.1* in blood cell lineages. CITE-seq data of healthy human bone marrow mononuclear cells (BMNC) and peripheral blood mononuclear cell (PMNC) populations. Data were retrieved from GEO (GSE139369) (Granja, Klemm et al. 2019). Myeloid cells include CD14^+^/CD16^+^ monocytes and granulocytes. T cells include T_CD4+_ and T_CD8+_. C) Gene track view of the *PU.1* locus showing an integrated single-cell ATAC-seq (scATAC-seq) and single cell RNA-seq (scRNA-seq) analysis of healthy human PBMCs. Data was from the 10k Multiome dataset (10x genomics). T cells includes T_CD4+_ and T_CD8+_.

### DNA methylation is anti-correlated with chromatin accessibility at the CREs in myeloid cells

DNA methylation is typically anti-correlated with gene transcription (Luo, Hajkova et al. 2018, Kribelbauer, Lu et al. 2020). Thus, we sought to examine methylation statuses at the CREs in blood cell lineages. To do so, we analyzed whole-genome bisulfite sequencing (WGBS) datasets of myeloid, B, T, and erythroid cells from a previously published atlas of human tissue methylation (GSE186458). Remarkably, we observed that the WGBS probes at all CREs were essentially unmethylated in myeloid cells shown by very low proportion of WGBS methylated probes (Figures 3A-B). Of note, some differential methylation was observed at several CREs in the three other lineages. While TL1, TL3 and TL4 were not methylated in all cell lineages, TL2, TL7, and TL8 were methylated in B, T, and erythroid cells but not myeloid cells. On the other hand, URE, TL5, hLZ12, PrPr, and the CpG island were specifically methylated in T cells, where *PU.1* expression was lowest among the cell lineages (Figures 2, 3A-B). Methylation at the CpG island is known to specifically correlate to gene silencing (Jones 2012, Luo, Hajkova et al. 2018). Taken together, our findings indicated that CRE unmethylation largely correlates with chromatin accessibility and high expression of *PU.1* in myeloid cells and that differential methylation statuses at the CREs link to decreasing *PU.1* expression in other blod lineages.

**Figure 3.**
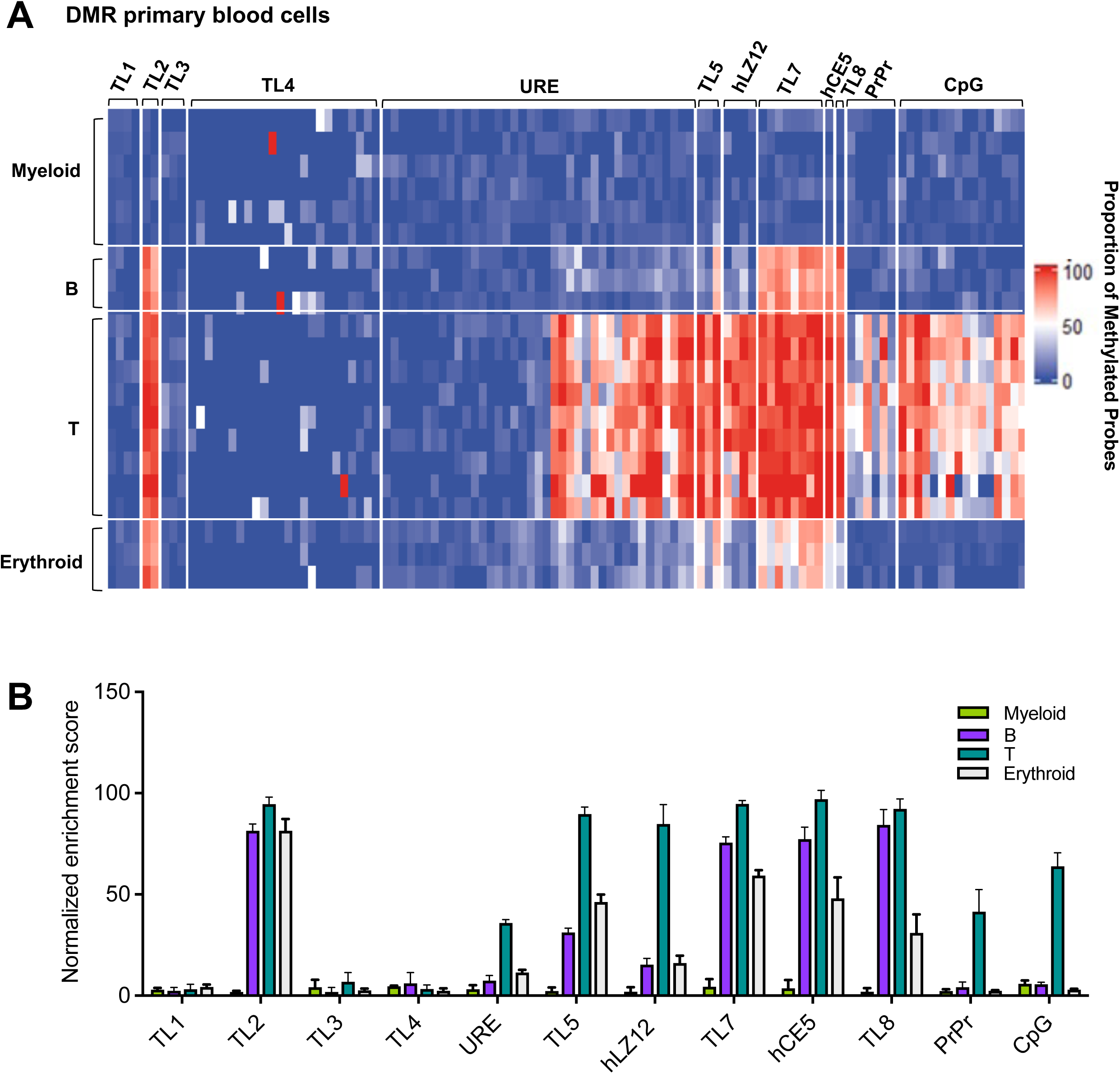
Differential DNA methylation statuses at the CREs. A-B) WGBS analysis of DNA methylation at the regions of interest within the *PU.1* locus in myeloid cells (monocytes and granulocytes), B, T, and erythroid cells. Dataset (GSE186458) was downloaded from GEO. A) Heatmap showing proportion of methylated probes at the regions of interest. B) Graphs of normalized methylation enrichment scores at CREs across blood cell lineages.

### An 8-kb CRE cluster at the upstream region of the *PU.1* gene is demarcated with differential H3K27Ac histone mark that reflect lineage-specific expression pattern

We next delineated the association between histone post-translational marks for enhancers and the CREs. Because the URE is the most experimentally well-defined enhancer in myeloid cells (Li, Okuno et al. 2001, Rosenbauer, Wagner et al. 2004, Okuno, Huang et al. 2005, Rosenbauer, Owens et al. 2006, Huang, Zhang et al. 2008, Zarnegar, Chen et al. 2010, Will, Vogler et al. 2015), we used this CRE as the benchmark to interrogate enhancer status at other *PU.1* CREs. Indeed, we revealed strong enrichment of the H3K27Ac active enhancer mark at a cluster of CREs spanning from the URE to the TL8 elements (Figure 4A). These enrichments also corresponded with chromatin opening (Figures 1B and 2C), lack of DNA methylation (Figure 3), and predominant *PU.1* expression (Figure 2) in myeloid cells. In B and erythroid cells, only weak H3K27Ac enrichment was noted at the URE. The putative CREs exhibited H3K27Ac enrichment in myeloid cells, though to a lesser extent compared to the URE (Figure 4A), suggesting their contributing roles as myeloid-specific enhancers to human *PU.1* expression. Besides H3K27Ac, we observed diffuse enrichment of H3K4Me1 throughout the *PU.1* locus as well as the upstream region that covers all the CREs. The strength of H3K4Me1 enrichment was reduced in a stepwise manner, ranging from myeloid cells, B, erythroid, and T cells in the same manner as seen in the expression data (Figure S1A). Likewise, a broad ∼5kb region spanning from the promoter towards *PU.1* exon 1 exhibited H3K4Me2 enrichment that also reduced going from myeloid to B, erythroid, and T cells (Figure S1B). Of note, three narrow upstream regions with H3K4Me2 enrichment containing exanimated CREs were seen in all four lineages (Figure S1B). These suggest that H3K4Me1 and H3K4Me2, which are implicated to indicate enhancer and transcriptional activities (Creyghton, Cheng et al. 2010, Pekowska, Benoukraf et al. 2011), could be contributing signatures but not specifically reflect enhancer activities at the CREs.

**Figure 4.**
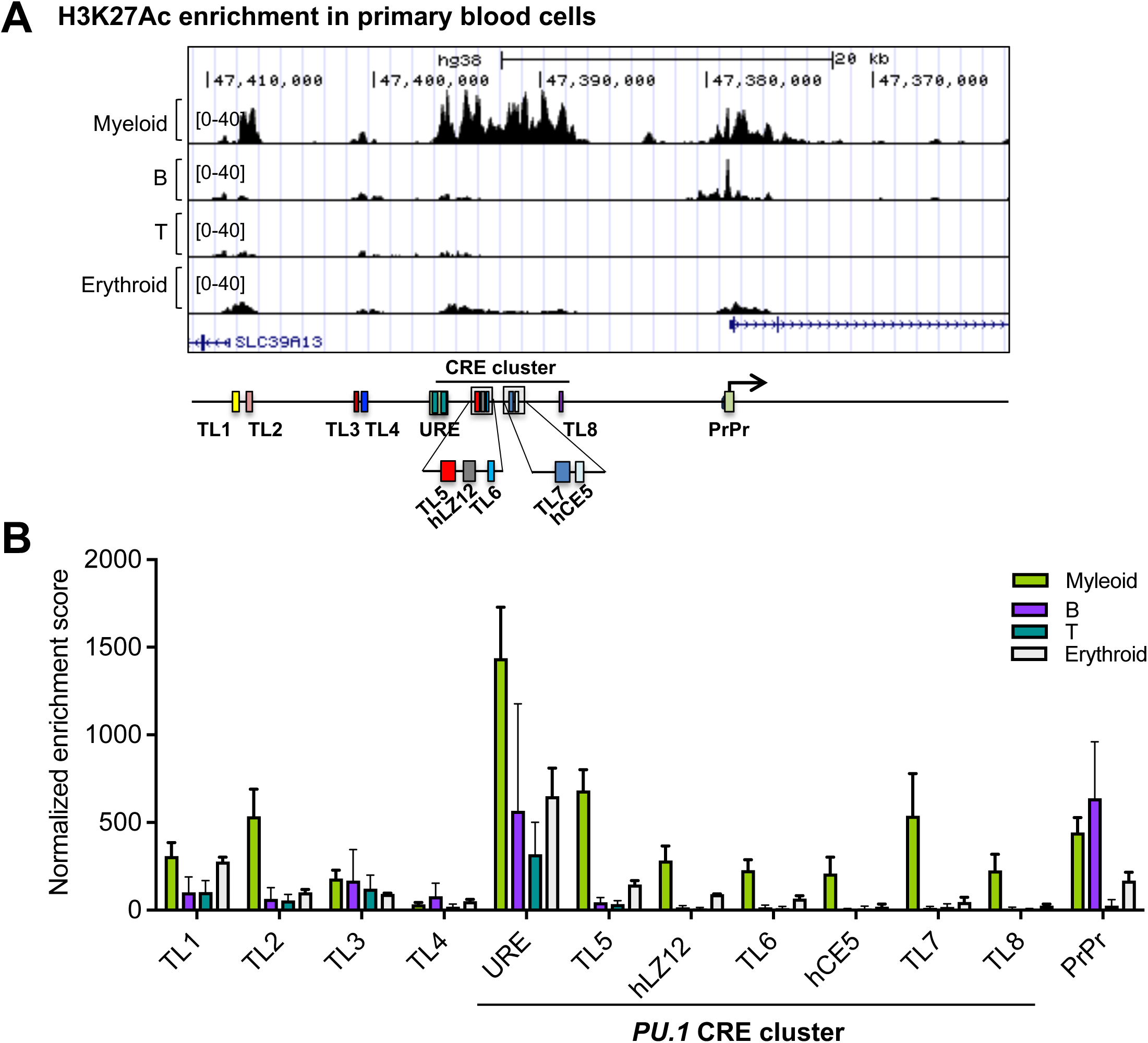
Enrichment of histone marks for enhancer and gene activation, see also Figure S1. H3K27Ac ChIP-seq tracks of four human blood cell lineages. Data were retrieved from ENCODE and GEO (GSE102584). A) Gene track view of the genomic region encompassing *PU.1* CREs. Shown are H3K27Ac ChIP-seq tracks of blood cell lineages. B) Bar graph of enrichment scores for H3K27Ac peaks at the CREs.

### *PU.1* CREs are not specifically demarcated with histone marks for silencer and gene repression in blood cell lineages

The URE is a bifunctional CRE, acting as an enhancer in myeloid cells and a silencer in T cells (Li, Okuno et al. 2001, Rosenbauer, Wagner et al. 2004, Okuno, Huang et al. 2005, Rosenbauer, Owens et al. 2006, Huang, Zhang et al. 2008, Zarnegar, Chen et al. 2010, Will, Vogler et al. 2015). Thus, we sought to examine histone marks that are implicated to demarcate silencers across blood cell lineages. To our surprise, we did not observe significant enrichment of H3K27Me3 repressive mark at the exanimated CREs in T cells or any other three lineages (Figure 5A). A similar finding was seen with H3K9Me3 (Figure 5B). In contrast, we found strong enrichments at other gene loci (Figures S2A-B), indicating that the lack of H3K27Me3 and H3K9Me3 at the *PU.1* locus was not an internal validity issue. Thus, the utility of H3K27Me3 and H3K9Me3 could be limited to inferring activity of broad genomic regions containing DNA elements in context-dependent manner rather than to pinpoint silencing activity of specific CREs at least in our exanimated blood cell lineages.

**Figure 5.**
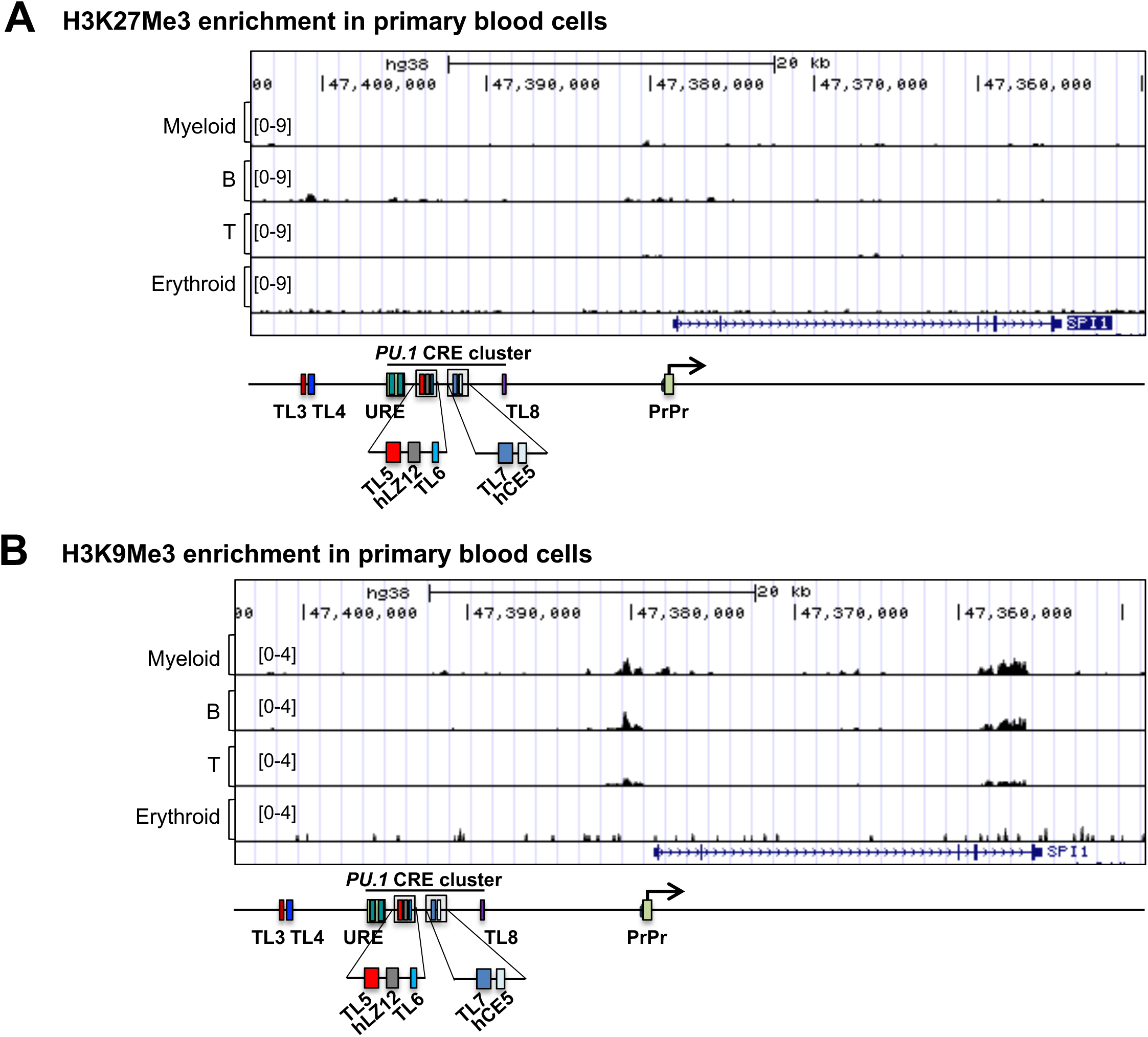
Histone marks for silencer and gene repression, see also Figure S2. A-B) Gene track view of the genomic region encompassing the *PU.1* locus and its upstream region. A) H3K27Me3 ChIP-seq tracks of four human blood cell lineages. Data were retrieved from ENCODE and GEO (GSE67783). B) H3K9Me3 ChIP-seq tracks of four human blood cell lineages. Data were retrieved from ENCODE and GEO (GSE12646).

### The CRE cluster is transcriptionally active and associated with *PU.1* autoregulation in myeloid cells

Active enhancers often exhibit transcriptional activities and are associated with trans-acting factor binding. For instance, histone 3 lysine 9 acetylation (H3K9Ac) and histone 3 lysine 4 trimethylation (H3K4Me3) are known to mark gene transcription and promoter activity (Pekowska, Benoukraf et al. 2011). Indeed, we revealed H3K9Ac and H3K4Me3 enrichment at the CRE cluster and in myeloid cells (Figures 6A-B and S3A-B). Such enrichments were found only at the URE and the PrPr in B cells (Figures 6A-B and S3A-B). On the other hand, only the URE exhibited these histone enrichments thought at a lesser level in T cells (Figures 6A-B and S3A-B). These results indicate that transcriptional activity at the CRE cluster is coupled with high *PU.1* expression in myeloid cells. To further elucidate this, we examined transcription factor binding at these CREs. Because PU.1 is known to autoregulate its own expression (Okuno, Huang et al. 2005, Huang, Zhang et al. 2008, Staber, Zhang et al. 2013), we focused our analysis on PU.1. Accordingly, we noticed PU.1 occupancy at the CRE cluster and the PrPr (Figures 6C and S3C). Intriguingly, even though *PU.1* expression is modest in B cells compared to myeloid cells (Figure 2), PU.1 occupancy at the URE was at comparable levels between myeloid and B cells. Notably, PU.1 occupancies at other CREs in the cluster were lower in B cells. This suggests that, in addition to regulation by activity at the URE, *PU.1* occupancies at other constituents of the CRE cluster link to superior *PU.1* autoregulation in myeloid cells.

**Figure 6.**
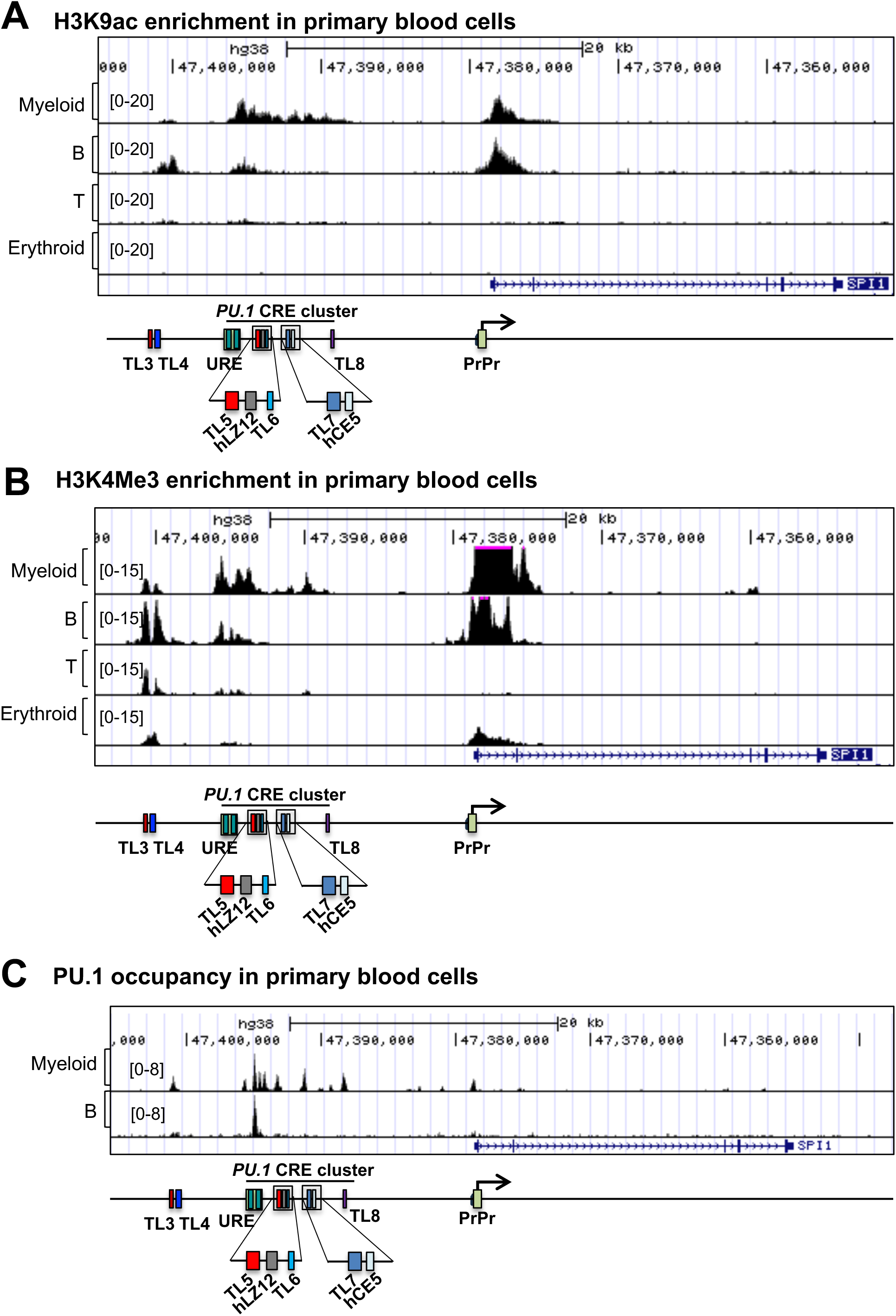
Enrichment of histone signatures for transcription and PU.1 occupancy at the CRE cluster, see also Figure S3. A-C) Gene track view of the genomic region encompassing the *PU.1* locus and its upstream region. A) H3K9Ac ChIP-seq tracks of four human blood cell lineages. Data were retrieved from ENCODE and GEO (GSE36985, GSE54634, GSE15735, GSE149295, GSE71422). B) H3K4Me3 ChIP-seq tracks of four human blood cell lineages. Data were retrieved from ENCODE and GEO (GSE102584). C) PU.1 ChIP-seq tracks of myeloid and B cells. Data were retrieved from ENCODE and GEO (GSE128834).

### Noncoding RNA transcripts are initiated at the CRE cluster in the upstream region of *PU.1* in myeloid cells

Transcription often occurs at active enhancers, giving rise to enhancer RNAs (eRNA). This includes 1d-eRNAs (long, polyadenylated and unidirectional transcription) and 2d-eRNAs (short, non-polyadenylated and bidirectional transcription) (Natoli and Andrau 2012, Li, Notani et al. 2016). We previously reported that the 1d-eRNA *LOUP* is originated from the URE and induces *PU.1* expression (Trinh, Ummarino et al. 2021). Therefore, we sought to comprehensively examine molecular indicators of transcriptional activities at the CRE cluster. Global Nuclear Run-On sequencing (GRO-seq) analysis (Core, Waterfall et al. 2008, Smale 2009) showed that, indeed, in addition to the gene body, nascent RNAs are present in the upstream region encompassing the CRE cluster, specifically in myeloid cells, with a superior frequency of transcription initiation (Figure 7A). To further elucidate this, we examined RNA polymerase II (Pol II) binding to chromatin and noted its occupancies at all the constituents of the CRE cluster and that its binding is more prominent at the URE (Figure 7B). Remarkably, RNAs were also detectable on the antisense strand with respect to the *PU.1* gene (Figure 7A). We further located transcription initiation sites within the CREs by inspecting the Cap Analysis Gene Expression sequencing (CAGE-seq) track from the FANTOM5 project (Kodzius, Kojima et al. 2006). Intriguingly, in addition to the CAGE-seq peak corresponding to 1d-eRNA *LOUP* initiation at the distal homology region (or H1) of the URE (Trinh, Ummarino et al. 2021), there are bidirectional CAGE-seq peaks within the proximal homology region (or H2) of the URE, suggesting the presence of a 2d-eRNA (Figure 7C). Moreover, we identified CAGE-seq peaks at TL5, hLZ12, TL7, and TL8, further indicating the presence of eRNAs as hallmarks for active enhancers (Figure 7C). Thus, the CRE cluster exhibits the molecular features of an enhancer cluster or a super-enhancer in myeloid cells.

**Figure 7.**
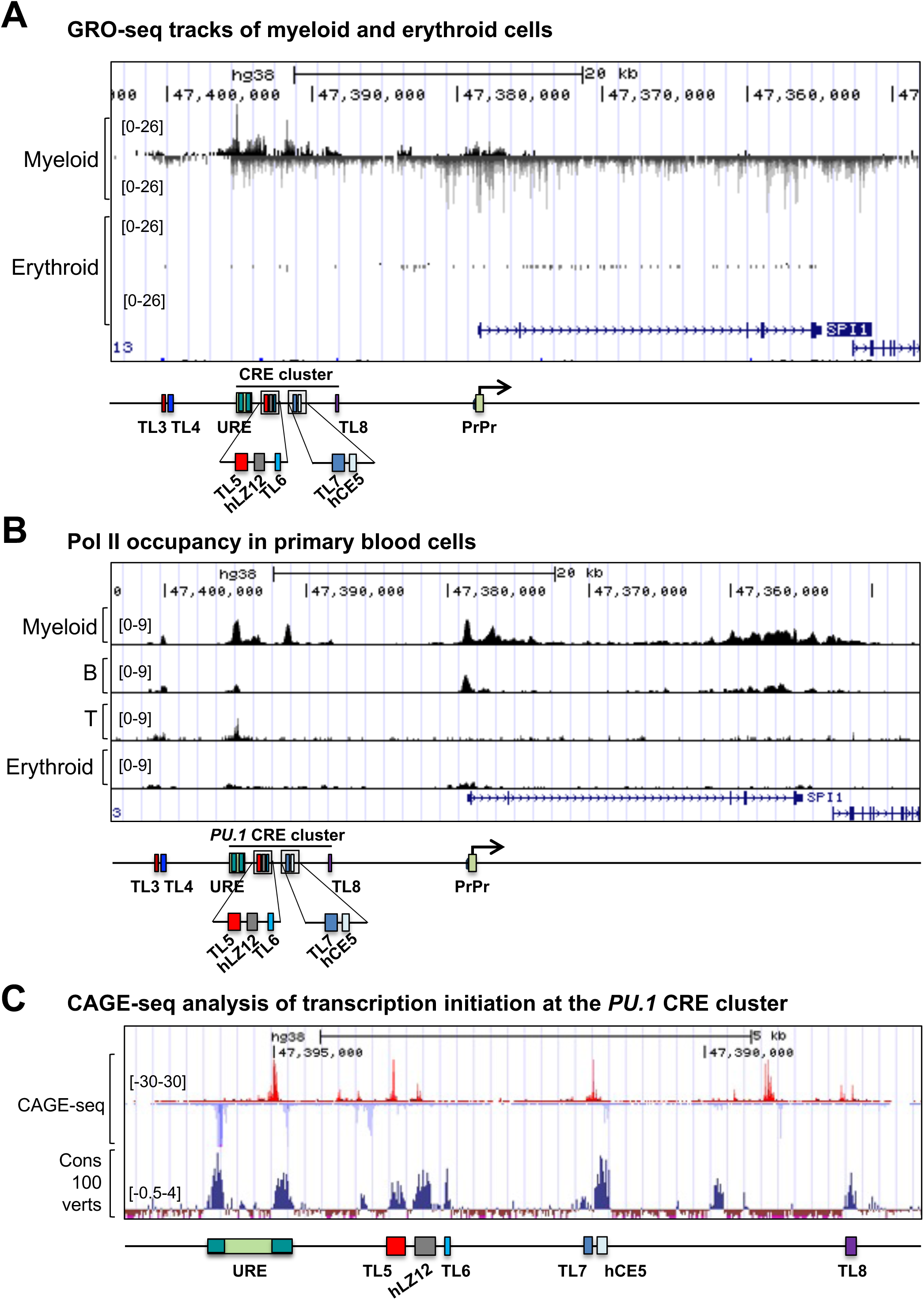
Noncoding transcriptional activities at the *PU.1* CRE cluster. A-C) Gene track view of the genomic region encompassing the *PU.1* locus and its upstream region. A) GRO-seq tracks of myeloid and erythroid cells. Data were retrieved from GEO (GSE136813, GSE102586). B) Pol II ChIP-seq tracks of four human blood cell lineages. Data were retrieved from ENCODE and GEO (GSE102586, GSE118305, GSE103477, GSE20040, GSE33281, GSE39538). C) CAGE-seq track from the FANTOM5 project.

### The presence of a lineage-specific CRE cluster-PrPr chromatin loop within the 35-kb CTCF-bound insulated neighborhood at the *PU.1* locus

Although the role of the URE-PrPr chromatin loop in *PU.1* induction in myeloid cells has been well described (Li, Okuno et al. 2001, Rosenbauer, Wagner et al. 2004, Okuno, Huang et al. 2005, Will, Vogler et al. 2015, Trinh, Ummarino et al. 2021), a comprehensive analysis of chromatin 3D architectures at the *PU.1* locus in human blood cell lineages has not been interrogated. To elucidate this, we examined Micro-C (or intact Mnase chromosomal conformation capture sequencing (Hi-C)) datasets that use micrococcal nuclease (MNase) digestion instead of traditional four or six-cutter restriction enzymes, enabling us to examine chromatin interactions at sub-kilobase resolutions (Hsieh, Cattoglio et al. 2020, Krietenstein, Abraham et al. 2020). To maximize the interaction map resolution, we called unbiased short-range loops at 1kb and 2kb resolutions simultaneously using HiCCUPS, which is capable of calling and merging loops at multiple resolutions together (Rao, Huntley et al. 2014). In all four blood cell lineages as well as a control non-hematopoietic cell lineage, we revealed a common loop formed between the TL1/2 region containing TL1 and TL2, which is accessible in all lineages (Figures 1C and 2C) and an uncharacterized region 4 kilobases downstream the *PU.1* promoter (P4K) (Figure 8A-E). To further characterize this loop, we inspected the occupancy of the DNA-binding protein CTCF, an important mediator of chromatin looping that represents the boundaries of potential chromatin loops. Indeed, we noted CTCF occupancy at these two elements (Figure 8A-D, bottom panel). This indicates that a CTCF-mediated chromatin loop forms a 35-kb insulated chromatin neighborhood at the *PU.1* locus in a lineage-independent manner. Interestingly, we noticed the presence of various lineage-specific chromatin interactions within this large chromatin loop in blood cells (Figure 8A-D). Of note, the interaction between the CRE cluster and the PrPr was detected in myeloid and B cells, of which the URE or the TL6 represent the interaction points (Figure 8A-B) corresponding to *PU.1* expression in these cells (Figure 2). In T cells, the CRE cluster shifted its docking from the PrPr to the P4K region (Figure 8C) in line with *PU.1* silence in these cells (Figure 2). In line with predominant chromatin accessibility at TL3 and TL4 in B and T cells (Figure 2C), TL3/4-P4K interaction was also noted in these cells (Figure 8B-C). In erythroid cells, where *PU.1* is minimally expressed (Figure 2), no CRE cluster-PrPr interaction was found. Instead, we noted the TL1/2-PrPr interaction (Figure 8D). In non-hematopoietic tissue (heart left ventricle), no chromatin interactions identified in blood lineages were present (Figure 8E). Taken together, these results indicate that chromatin interactions between CREs and the PU.1 promoter in 3D space are a major contributing factor to lineage-specific *PU.1* expression.

**Figure 8.**
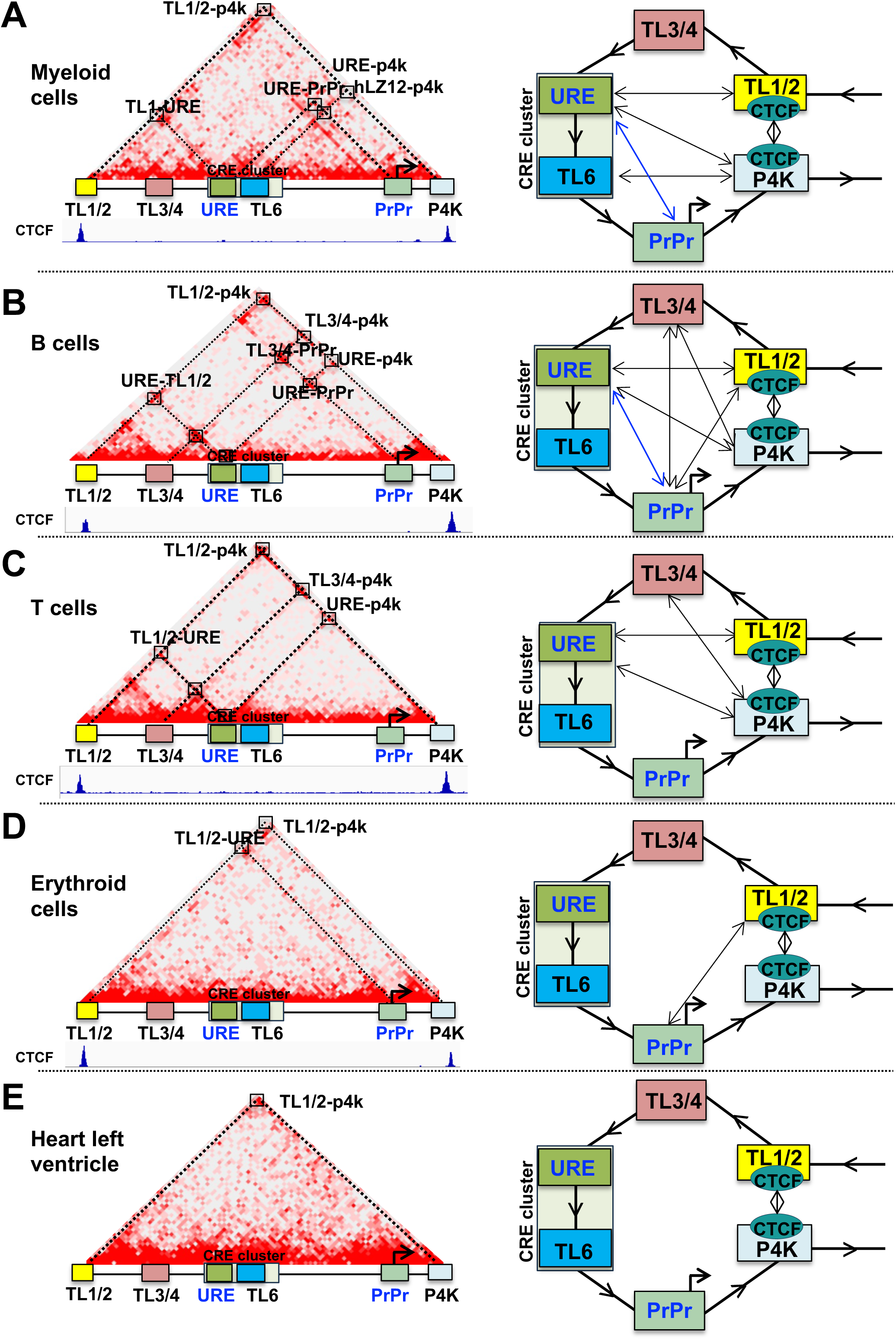
Chromatin architecture dynamics at the *PU.1* locus in cell lineages. Hi-C contact map (left) and schematics for chromatin interactions (right) of A) myeloid cells (CD14+ monocytes, ENCODE: ENCSR236EYO)), B) B cells (ENCODE: ENCSR847RHU), C) T cells (CD4+ T cells, ENCODE: ENCSR458VJJ), D) erythroid cells (erythroid precursors, GEO: GSM6616197), and E) heart left ventricle (ENCODE: ENCSR998LLZ). Corresponding genomic locations with CREs, and CTCF ChIP-seq tracks are shown underneath each contact maps (data were retrieved from ENCODE and GEO (GSE36985 and GSE67783)). In the schematics, the CRE cluster with interaction points, URE and TL6 are shown. Arrows represent chromatin direction starting from upstream region toward the PU.1 gene body. Two-direction arrows depict chromatin interactions identified by HiCCUPS.

In summary, we have identified novel CREs at the upstream regions of the human *PU.1* locus and demonstrated their role in *PU.1* regulation via chromatin architecture. A subset of CREs localize within a cluster that exhibits molecular features and chromatin signatures of a super-enhancer in myeloid cells. The CREs involved in lineage-specific chromatin interactions within a universal and insulated neighborhood (Figure 9). Our findings raise the possibility that chromatin signatures and architectures acts in concert to regulate gene expression providing new insights on the bifunctional role of CREs.

**Figure 9.**
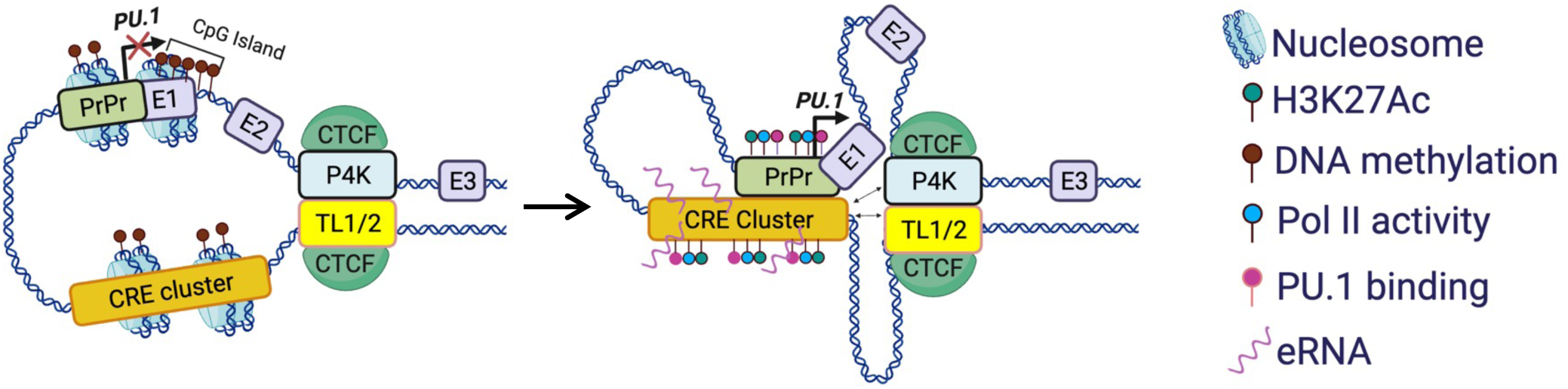
A model of chromatin structure and 3D architecture dynamics in *PU.1* long-range transcription regulation. The common 35-kb CTCF-bound insulated neighborhood with TL1/2 and P4K boundaries spanning from 29 kilobase upstream and 4 kilobases downstream of the *PU.1* transcription start site. In the resting state, the CRE cluster and the PrPr are heavily methylated and inaccessible (left panel). In the highly active *PU.1* transcription state (middle panel), the CRE cluster exhibits super-enhancer activities with unmethylated DNA, chromatin opening, Pol II activity, eRNA production, PU.1 occupancies, and 3D chromatin remodeling coupled with functional CRE cluster-promoter interactions. Shown are involving *PU.1* exons (E1, E2, E3). Two-way arrows represent additional chromatin interactions.

## DISCUSSION

In this study, we discovered a myeloid-specific CRE cluster at the upstream region of the *PU.1* gene exhibiting molecular signatures that help explain the dynamic expression pattern of *PU.1* in blood cell lineages. Our study revealed several important findings. We demonstrated that chromatin accessibility and unmethylation status at certain constituents of the CRE cluster is correlated with the distinct expression pattern of *PU.1* in four major blood cell lineages. The CRE cluster is demarcated with differential H3K27Ac histone marks reflecting enhancer activities in myeloid cells but not specifically marked with histone modifications reported to be molecular indicators of silencer activities. We also revealed transcriptional activities and the presence of eRNAs at several constituents of the CRE cluster indicating its role as a *PU.1* super-enhancer in myeloid cells. Furthermore, we identified that PU.1 occupancies at the CRE cluster associates with *PU.1* autoregulation in myeloid cells. Finally, we revealed the presence of a hematopoietic-specific chromatin interactions that are wrapped within the CTCF-mediated non-lineage specific chromatin loop at the *PU.1* locus. These findings provide molecular insights into *PU.1* regulation via 3D chromatin architecture and link chromatin signatures to the dynamic expression pattern *of PU.1* in blood cell lineages.

Our finding of a myeloid-specific CRE cluster is intriguing. Based on sequence conservation and chromatin accessibility, identified eleven discreet CREs, including the URE. Not included in this count are two previously reported elements that were excluded as they did not pass our selection criteria. These include the murine CE4A (-10 kb) (Zarnegar, Chen et al. 2010) which exhibited sequence conservation but no chromatin accessibility in none of the four blood cell lineages, and the human -14.4 kb element (Dluhosova, Curik et al. 2014) that exhibits no sequence conservation among species. Among the CREs, we noted three was blood ubiquitous exhibiting similar chromatin accessibility and DNA unmethylation patterns across all four blood lineages. The remaining elements localize within an 8 kb-wide CRE cluster spanning from the URE to the TL8 and exhibit chromatin and epigenetic signatures for myeloid-specific enhancers shown by chromatin opening, lack of DNA methylation, and the presence of H3K27Ac enrichment specifically in myeloid cells. In examining further molecular indicators of active enhancers, we noted the presence of 1d-eRNAs and 2d-eRNAs, important hallmarks of active enhancers at several CRE constituents. In supporting the autoregulatory role of PU.1 in myeloid cells, we revealed PU.1 occupancies at the CRE cluster. This superior *PU.1* autoregulation in myeloid cells is likely an attribute of the CRE constituents other than the well-known URE. This is because even though *PU.1* expression is modest in B cells in comparing to myeloid cells, its occupancy at the URE was noted at comparable levels between myeloid and B cells. In contrast, PU.1 occupancies were much less at other CREs. Experimental examination of individual CREs within the *PU.1* CRE cluster is warranted for a further detailed understanding of their mechanistic actions and is being actively investigated.

How the same CRE can exhibit bifunctionality as enhancer and silencer and what are molecular signatures for silencers remain outstanding research questions. Indeed, among the *PU.1* CREs, the URE has been shown to demonstrate this bifunctionality in prior ablation studies as both an enhancer in myeloid cells and a silencer in T cells (Rosenbauer, Wagner et al. 2004, Rosenbauer, Owens et al. 2006, Will, Vogler et al. 2015, Willcockson, Taylor et al. 2019). As comprehensively examined in this study, we identified a repertoire of CREs that exhibit molecular signatures for enhancers in myeloid cells. These include, unmethylation, H3K27Ac enrichment, the presence of eRNAs, as well as trans-acting factor occupancies including PU.1 and polymerase II at the CREs. With regards to molecular signatures for *PU.1* silencers in T cells, however, we could not observe specific enrichment of the repressive mark H3K27Me3 (Young, Willson et al. 2011, Huang, Petrykowska et al. 2019) and the heterochromatin mark H3K9Me3 (Nicetto, Donahue et al. 2019) neither at the URE nor any *PU.1* CRE constituents. This is understandable because, indeed, only a small portion of silencers exhibits H3K27Me3 and H3K9Me3 enrichment (Young, Willson et al. 2011, Huang, Petrykowska et al. 2019, Doni Jayavelu, Jajodia et al. 2020). Thus, these histone marks seem limited to inferring specific gene-silencer characteristics and not defining broader silencer regions, which speaks to the challenges of defining enhancing and silencing CREs using existing high-throughput assays. Further studies into histone modification marks of CRE bifunctionality especially chromatin signatures for silencers are needed not only for *PU.1* CREs but also silencers in general.

Our finding of lineage-specific chromatin interactions involving the *PU.1* CREs within a common 35-kb insulated neighborhood mediated by CTCF provides two important insights into CRE-mediated long-range transcription regulation. First, the insulated neighborhood could form a chromatin territory for functional CRE-PrPr interactions to occur. Indeed, this chromatin loop is located within a bigger chromatin loop, the 75-kb SubTAD (Schuetzmann, Walter et al. 2018). Second, lineage-specific chromatin interactions within this insulated neighborhood could link to differential *PU.1* expression in blood lineages. These include functional interactions of the CRE cluster as a super-enhancer to the *PU.1* promoter in myeloid and B cells and the shifting of CRE cluster interaction from the promoter to the P4K region in T cells. Previous studies using 3C assays also detected the shift to the AsPr further downstream of the P4K region in T cells (Ebralidze, Guibal et al. 2008, van der Kouwe, Heller et al. 2021). Although we were unable to detect this interaction in Micro-C data, it is possible that this interaction is transient and versatile that the technology is unable to capture. Also, with the limitations in current chromatin conformation capture technologies, it is not yet feasible to discern whether multiple PU.1 CRE interactions within the insulated neighborhood occurred sequentially or simultaneously in individual cells. Nevertheless, our study supports the model that dynamic and alternative CRE dockings in different cell lineages define their functions as enhancers or silencers. This provides a divergent observation from the model that the same CRE could function as an enhancer and silencer by utilizing the same CRE-promoter chromatin loop (Huang, Petrykowska et al. 2019), at least for the *PU.1* locus.

In summary, we performed an unbiased and holistic analysis on public high-throughput sequencing datasets to identify novel CREs at the upstream regions of the human *PU.1* locus. A *PU.1* CRE cluster with known and unknown CREs exhibits chromatin signatures of a myeloid-specific super-enhancer in myeloid cells. We also revealed a universal and insulated neighborhood that contains the CRE cluster, forming the chromatin territory for lineage-specific chromatin interactions that involves the CREs. This forms dynamic 3D chromatin architectures linked to specific *PU.1* expression patterns in blood cell lineages. Our findings set lights on the bifunctional nature of certain CREs on chromatin regulation in a context-dependent manner, thus providing mechanistic insights into the epigenetic and chromatin regulation of *PU.1*. Further experimental studies will examine the interplay between *PU.1* CREs, chromatin structure, and 3D architecture in malignant hematopoiesis.

## ACKNOWLEDGEMENTS

This work was supported by the following grants and awards. NCI K01 CA222707, Grant # 134088-IRG-19-143-33-IRG from the American Cancer Society, and state funding within the UVA Comprehensive Cancer Center and UVA School of Medicine to BQT; NCI R35 CA197697, R01DK103858, W81XWH-15-1-0161, and P01HL131477-01A1 to DGT, NIH R01 HL130550, and R01 HL149667 to ANG, VCU Quest Award to TND. We thank Tuan Nguyen, UVA Research Computing staffs, and members of the Trinh lab for assistance and helpful suggestions.

## SUPPLEMENTAL FIGURE LEGEND

**Figure S1.**
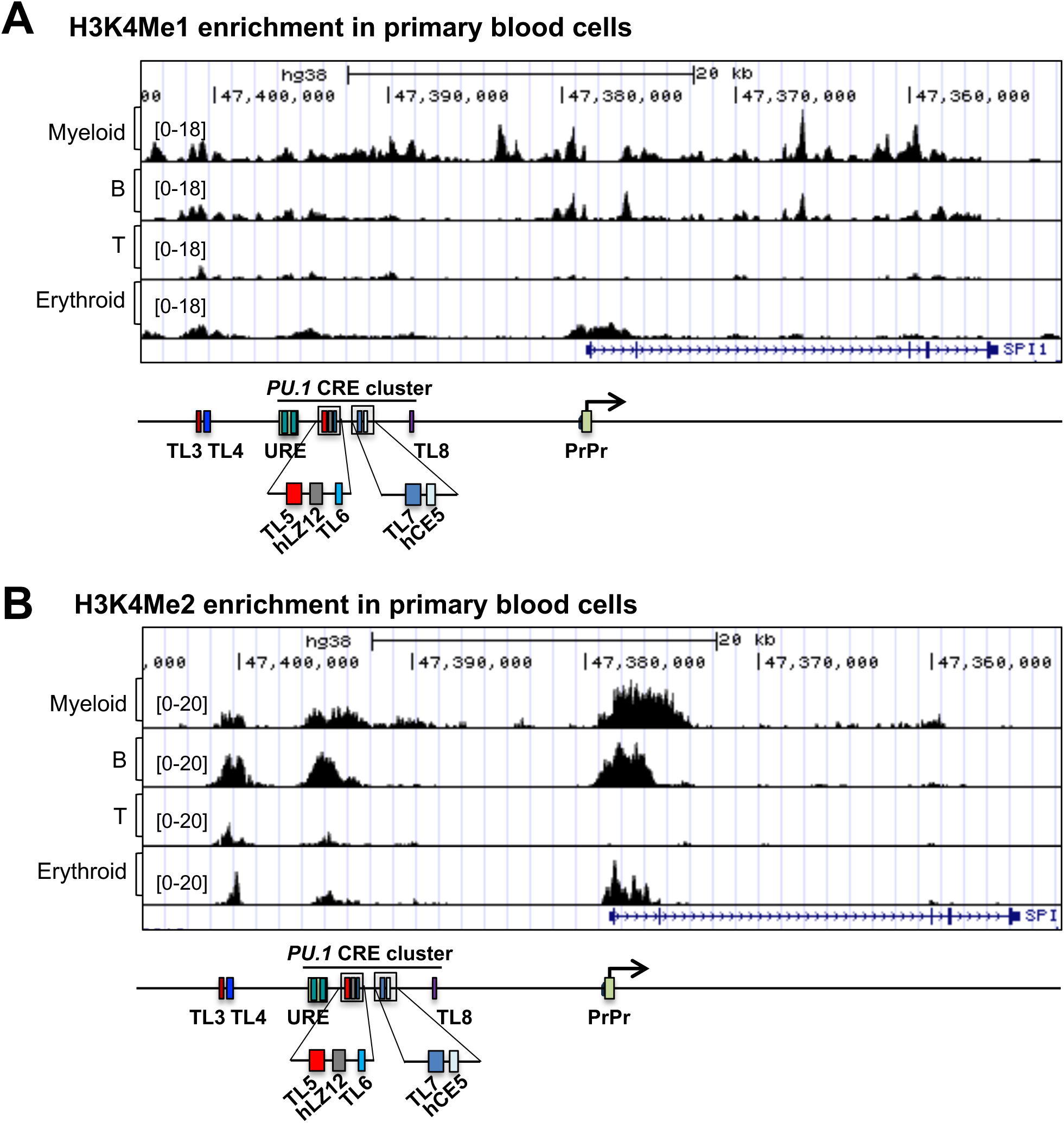
Histone marks for enhancer and gene activation, Related to Figure 4. A-B) Gene track view of the genomic region encompassing *PU.1* CREs. Shown are A) H3K4Me1 and B) H3K4Me2 ChIP-seq tracks of human blood cell lineages. Data were retrieved from ENCODE and GEO (GSE102584 and GSE73214)

**Figure S2.**
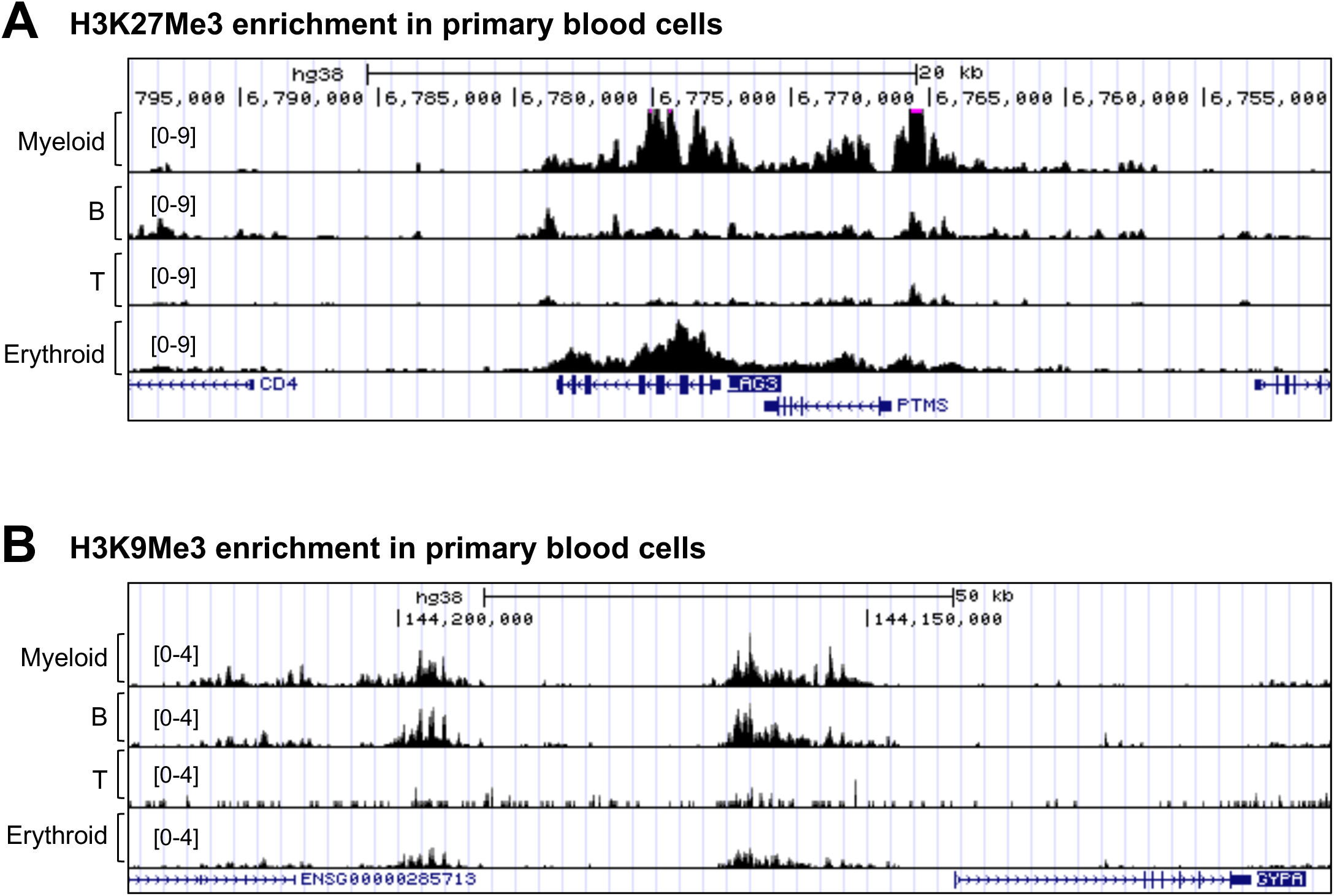
Histone marks for silencer and gene repression, Related to Figure 5. A) Gene track view of the genomic region encompassing the *LAG3* and *PRMS* loci. H3K27Me3 ChIP-seq tracks of four human blood cell lineages. Data were retrieved from ENCODE and GEO (GSE67783). B) Gene track view of the genomic region encompassing the *GYPA* locus. H3K9Me3 ChIP-seq track of four human blood cell lineages. Data were retrieved from ENCODE and GEO (GSE12646).

**Figure S3.**
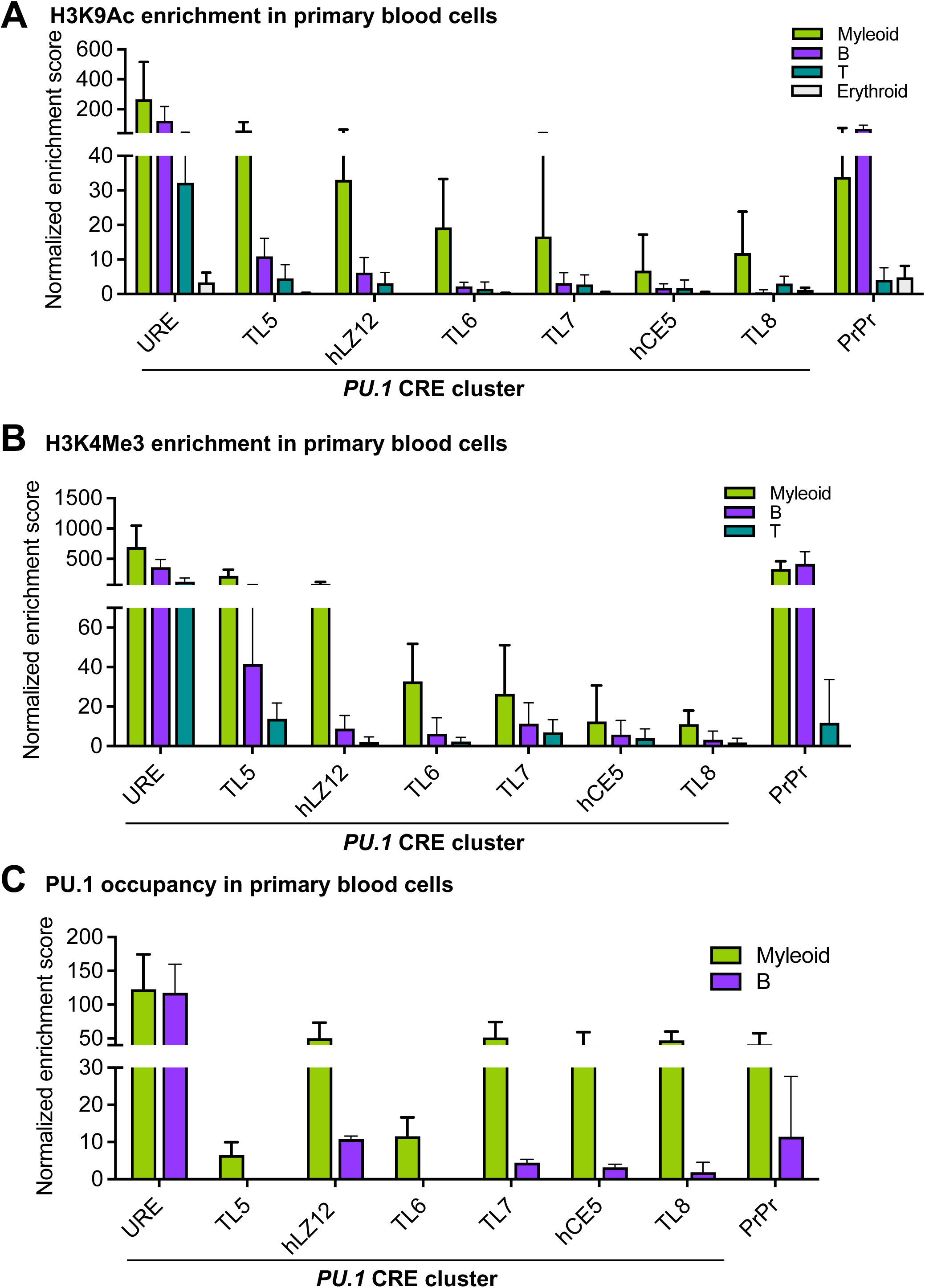
Enrichment of transcription histone signatures and PU.1 occupancy at the CRE cluster, Related to Figure 6. A-C) Bar graphs of enrichment scores for H3K9Ac (A), H3K4Me3 (B), and PU.1 (C) ChIP-seq peaks at the CREs. See figure 6 for details.

